# On interactive spatial visualisation of pathogenicity predictions

**DOI:** 10.1101/2025.03.19.644131

**Authors:** Junjie Xu, Aaron S. Kovacs, Stephanie Portelli, Ashar J. Malik, David B. Ascher

**Author notes:** Corresponding authors: Ashar J. Malik & David B. Ascher.

## Abstract

The past few years have revolutionised the utility of protein structural information in assessing disease risk. Such efforts have been pioneered by the DeepMind and Meta teams, who initially developed accurate protein prediction tools AlphaFold and ESMFold, respectively. More recently, both have been readapted to create AlphaMissene and ESM1b, through which, the pathogenicity potential of all possible missense mutations within the human proteome has been predicted. Similarly, VESPA was created as an alignment-agnostic predictor, which infers conservation from protein-language model embeddings to assign an effect for each variant. To facilitate their utility, we present a unified dashboard, MissenseViewer, which permits direct predictor comparison, and visualisation within the protein 3-dimensional structural context. Our interactive resource is freely available at: https://biosig.lab.uq.edu.au/missenseviewer/.

## Introduction

The once-elusive promise of precision medicine has today become more practically achievable. Significant breakthroughs, both in terms of quality, and cost of sequencing genomic data, has allowed clinicians to access a comprehensive snapshot of their patient genomes. Clinical annotations of missense mutations, however, have struggled to keep up, uncovering an opposite problem: the availability of comprehensive, but incomprehensible data. Computational prediction tools^1-4^ have sought to fill this data knowledge gap, by relying on conservation trends across species to assess the importance of specific amino acids to protein structure and function. Relying on limited information, these tools have exhibited minimal success in accurately detecting disease.

We have previously shown that assessing missense mutations within their 3-dimensional (3D) protein context offers invaluable information for detecting both drug resistance^5-7^ and disease risk^8,9^. Recent advances in protein structure prediction have made this approach more feasible, as the accurate structures of most of the human proteome are now accessible through tools like AlphaFold2^10^ and ESMFold^11^. Such tools dissect critical information from queries and homologs and iteratively optimise intermediary predicted structures to produce a final model. Similar approaches have been more recently used to develop AlphaMissense^12^ and ESM1b^13^, respectively, to predict the effect of missense mutations.

Both tools exhibited high robustness in detecting pathogenic variants. ESM1b reported similar performances on their sensitivity (81%) and specificity (82%)^13^ meaning that their predictor does not have an underlying bias in assigning one phenotype over another. AlphaMissense^12^ outperformed ESM1b on various clinical datasets, as reported through the area under the Receiver Operating Characteristic (auROC) curve, a metric which combines sensitivity and specificity. In practice, both methods predict a specific score for each missense mutation, which, due to the different architectures used, lie across different numerical scales: “0” to “1” in AlphaMissense, and “-30.945” to “16.896” in ESM1b. These scores are subsequently used to designate the mutant to a benign or pathogenic class, and an additional ‘ambiguous’ class in AlphaMissense.

Notably, both these methods have been successful in extraction of evolutionary patterns which highlights regions of functional importance across homologs. In both cases, high dimensional embeddings are used to determine effects of variants at each site. While this approach is rightfully deemed revolutionary, the team behind VESPA^14^ (variant effect score prediction without alignments), have competitively predicted variant effects using a different framework. This approach proved comparable in performance (Spearman correlation coefficient, *r*) to other state-of-the-art approaches, including the ESM1b precursor, ESM1v.

The prominence of these methods towards precision medicine has been affirmed not only through the robustness with which they predict variant effects, but also through their initiatives in scaling these predictions. Both AlphaMissense and ESM1b have predicted phenotypes of all possible missense mutations across the human proteome and made them publicly available, whereas pathogenicity inference from VESPA can be easily made through its predictions of variant fitness. Given the high performance achieved by these methods, they can not only directly guide further precision medicine efforts but also permit the knowledge-driven development of novel therapeutics.

However, the availability of numerical predictions without their 3D structural context offers a rather restricted representation and is insufficient to appropriately analyse consequences of missense mutations. This is because upon folding, proteins can adopt different folds where subtle changes through missense mutations can be tolerated at varying extents^15^. For example, the substitution of a residue at the protein surface might be more tolerated compared to one at the protein core. Specific protein classes, like tyrosine kinases, are also capable of exploring significantly different conformations under different binding conditions^16^, meaning that the effect of missense mutations can vary depending on the state, often controlled by interplay of a different subset of residues independent from the mutation site. Including structural context offers knowledge-driven ways to comprehensively interpret predictions from all three models, particularly when comparing predictions in buried and exposed regions, ordered with disordered parts of the structure, and different proximities to functional sites (Figures 1-4).

**Figure 1.**
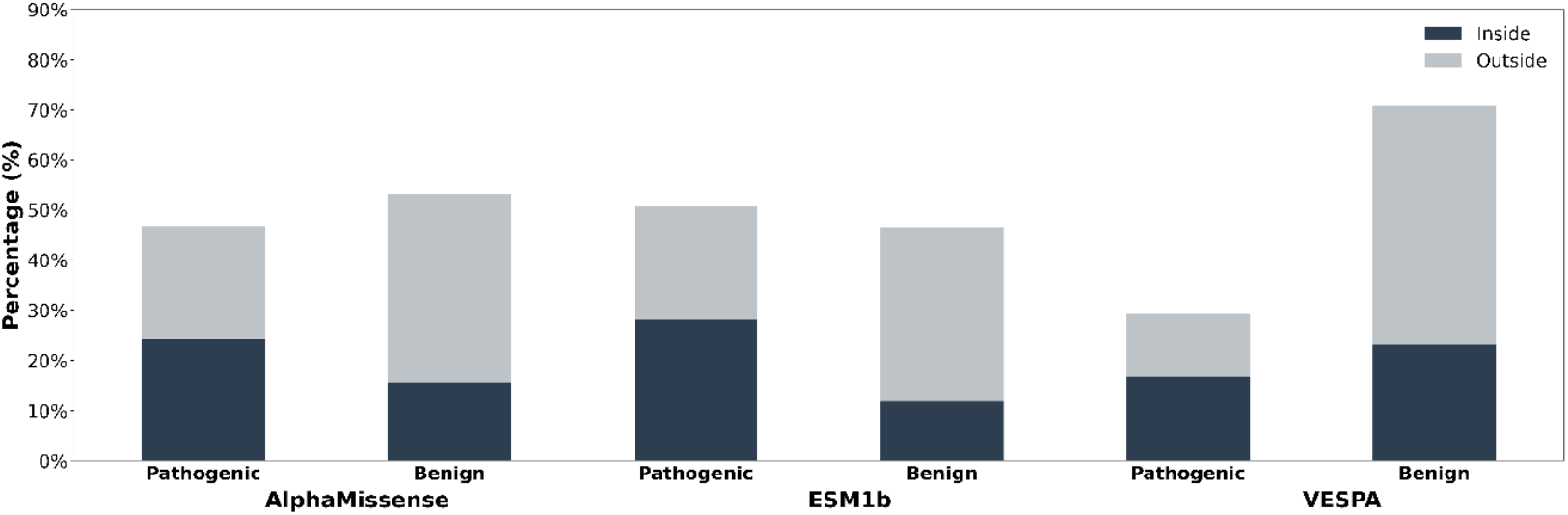
Variations classified as pathogenic and benign by AlphaMissense, ESM1b and VESPA. Fractions of each class within (blue) and outside Pfam domains (grey) is also shown.

**Figure 2.**
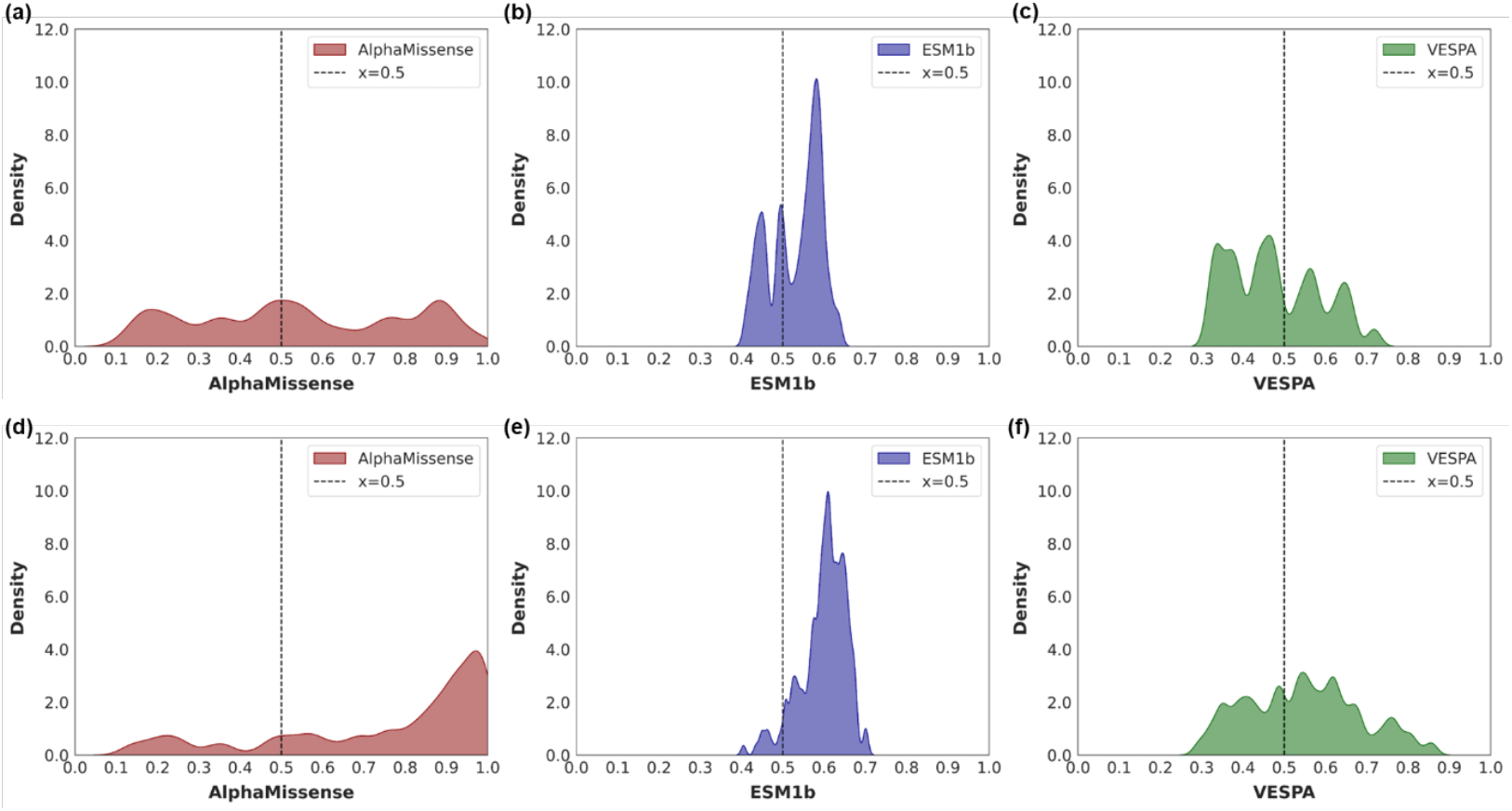
Distribution of predictions for variation within protein TP53 (Uniprot accession P04637) tetrametization domain (a-c) and DNA binding domain (d-f). Within the tetramerization domain (a-c), AlphaMissense and VESPA show a uniform spread of benign (≤ 0.5) and pathogenic (> 0.5) predictions. ESM1b, on the other hand predicted a greater number of site variations as pathogenic. When considering the DNA binding domain (d-f), all three methods predict most sites as pathogenic (> 0.5).

**Figure 3.**
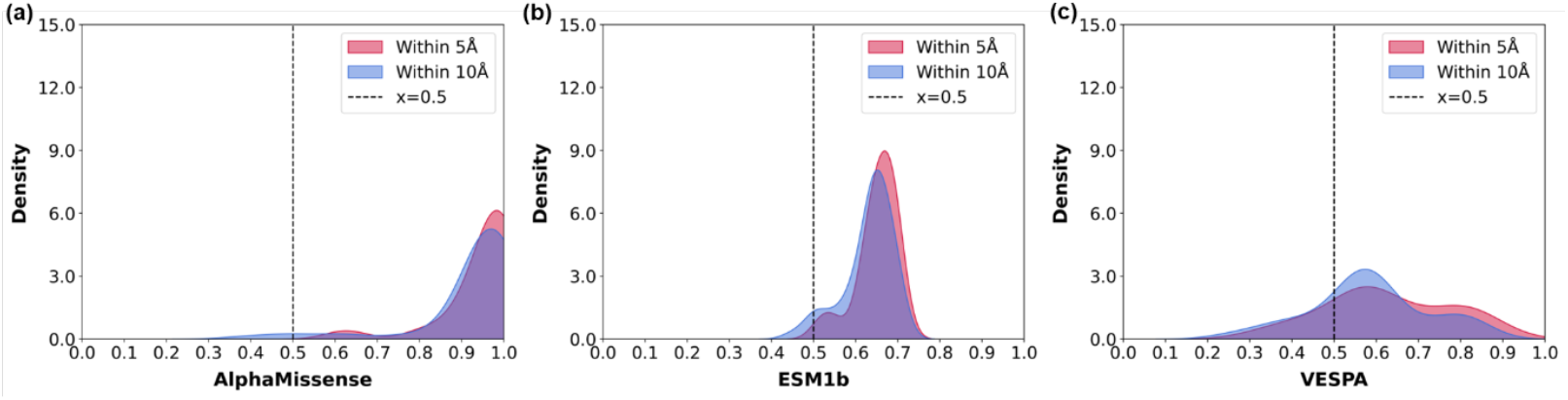
Distribution of prediction score at positions close (within 5 Å) or distant (within 10 Å) to the CDK2 ATP binding site (PDB ID: 2CHH). All three methods highlight higher rates of predicted pathogenic variation (> 0.5) at these sites, with slightly more prominent distributions for residues within 5 Å (red) compared to those within 10 Å (blue).

**Figure 4.**
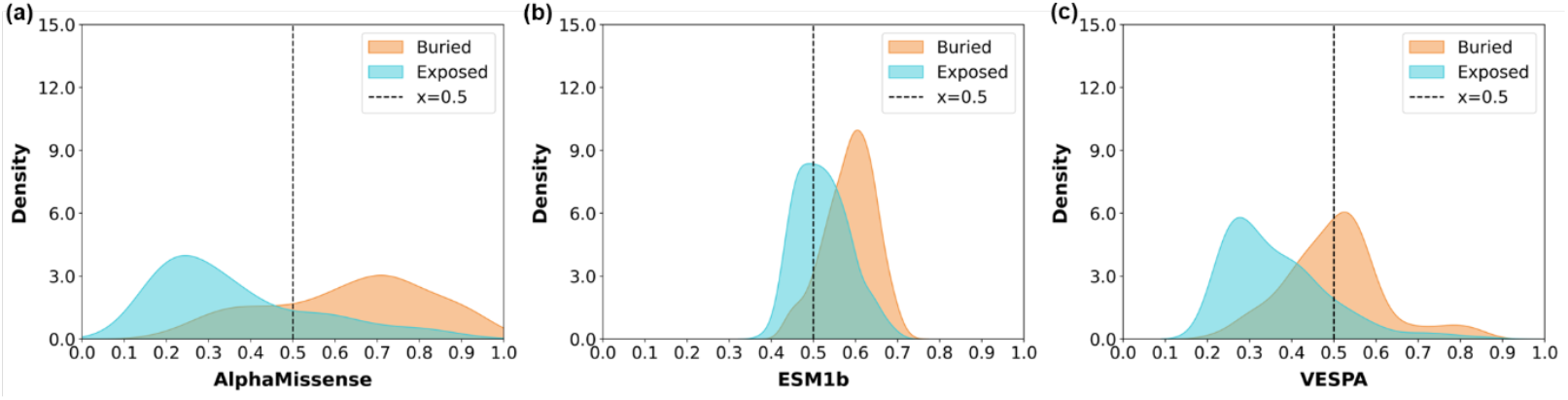
Distribution of prediction scores at buried (surface accessibility < 30%) and exposed (surface accessibility > 30%) positions in the cytochrome P450 1A1 protein (PDB ID: 6O5Y). All three methods highlight higher rates of predicted pathogenic variation (> 0.5) at buried sites (orange), compared to exposed sites (teal).

To address this, we have developed MissenseViewer, a web resource which complements predictions provided by AlphaMissense, ESM1b and VESPA with protein structural information. Through this interactive resource, users are not only able to access predictions across the human proteome but also visualise functional patterns on their respective protein 3D structures. We have made this freely available at: https://biosig.lab.uq.edu.au/missenseviewer/.

## Methods

AlphaMissense and ESM1b scores were collated from their repositories as cited in their publications, which identified a total of 19,838 protein isoforms which were common across the methods. Uniprot accessions of this isoform subset were used to recover respective fasta sequences from the ESM1b dataset which were then submitted to VESPA to generate effect scores per site, per mutation.

ESM1b scores, which ranged between “-30.945” to “16.896” were inverted and scaled to the range of scores covered by AlphaMissense and VESPA: between “0” and “1”, for appropriate comparison. In doing so, the cutoff between effect and no-effect (ESM1b score = -7.5) was mapped to the middle of the normalised range, 0.5. The scores from all unique mutations at the same site were then averaged to arrive at a per site prediction score for every predictor, which will henceforth be referred to as “prediction score”. For AlphaMissense scores, which traditionally have two cutoffs separating mutations into benign, ambiguous and pathogenic, we considered one mid-point cutoff score to distinguish mutations into two classes: those with values ≤0.5 were regarded as benign and those with values >0.5 as pathogenic. This was primarily done to determine general agreement between the three methods, while exact averages have been used in all other instances.

EBI-SIFTS^17,18^ was next used to retrieve all RCSB PDB structures for the previously-identified Uniprot accessions. The average susceptibility score at each position was multiplied by “100” and mapped onto the protein structure B-factor column to permit visualisation of effects on protein structures. AlphaFold structures were downloaded from the AlphaFold database hosted on EBI. For ligand association, HET codes were used to identify non-amino acid molecules and biopython^19^ was used to determine proximal residues in protein at 5 Å and 10 Å distance cutoffs. Solvent accessibility was determined using Naccess^20^ and a threshold of 30% was used to determined exposed and buried residues.

ClinVar variants were downloaded from the NCBI website (accessed on 07/10/2023). To map the scores of the models to these variants, the genome location and nucleotide substitution were extracted from the main ClinVar file. Running these through the Ensembl tool, VEP^21^, produced the transcript, residue site and resulting amino acid of each variant. This information could then be used to map the ClinVar variant to its position within the canonical Uniprot protein and therefore also to the AlphaMissense and ESM1b scores.

The webserver utilized flask on the backend and standard HTML and javascript on the frontend. Mol*^22^ was used to render protein structures and RCSB saguaro^23^ was used to display features.

## Results

In comparing the level of congruency between the three methods for each site across the human proteome (Table 1), we observed a high level of agreement between VESPA, which predicts variant effects, with AlphaMissense and ESM1b, which are trained to predict pathogenicity. Due to this, it can be assumed that for the human proteome, the predicted effect from VESPA can be considered to have pathogenic potential.

**Table 1:** Percentage of sites across the human proteome where the prediction of the three methods congruently give the same classification.

| Method | AlphaMissense | ESM1b | VESPA |
| --- | --- | --- | --- |
| AlphaMissense | 100 | 77.20 | 74.84 |
| ESM1b | 77.20 | 100 | 69.11 |
| VESPA | 74.84 | 69.11 | 100 |

We have also integrated Pfam domain information within MissenseViewer, which highlights domains and regions of evolutionary and hence, functional, importance across the proteins. We have compared the proportions of predicted pathogenic and benign variation as they localised either inside or outside Pfam domains and found that variation was more consistently predicted benign in protein regions outside of Pfam domains (Figure 1). This suggests that all tools can capture evolutionary patterns to predict final phenotypes. Notably, however, while AlphaMissense and ESM1b have approximately equivalent proportions of predicted pathogenic and benign variants, it is also observed that VESPA more often predicted variants as benign compared to pathogenic, within our protein dataset.

To further illustrate utility of these predictions, we next looked at predicted scores of variation at the tetramerization (Figure 2a-c) and DNA binding domains (Figure 2d-f) within TP53 (Uniprot accession: P04637). Notably, it was observed that higher proportions of mutations lining the DNA binding domain were predicted pathogenic across the three methods, compared to those in the tetramerization domain. Notably, in both regions, ESM1b predicted a larger proportion of mutations to be pathogenic (>0.5), suggesting it might be more sensitive to functional regions. In line with this, Figure 1 shows how fluctuations in pathogenic and benign proportions for ESM1b predictions are more prominently reliant on Pfam domains than the other predictors tested.

Next, we explored how the predictions from AlphaMissense, ESM1b and VESPA range according to distance to specific functional sites. Using the kinase CDK2 (Uniprot accession: P24941) as a case study we adopted a site-specific analysis looking at score distributions for residues within 5 and 10 Å from bound ATP, using an ATP-bound structure (PDB ID: 2CHH). These distances were used to highlight regions within the kinase which are directly (5 Å) and indirectly (10 Å) interacting with ligand ATP, thereby possibly affecting its function. Interestingly, all three predictors predicted these variants as pathogenic (> 0.5), where in all cases, residues within 5 Å had consistently slightly higher values. Interestingly, considering values obtained, AlphaMissense (Figure 3a) was observed to have the highest values, were predictions peaked at around 0.98, followed by ESM1b (Figure 3b; 0.68) and VESPA (Figure 3c; 0.58). This suggests that AlphaMissense better captures functional properties of a protein, as exemplified through the kinase ATP-binding site. Interestingly, VESPA predicts effects in a less binary manner, highlighting that accounting for protein fitness correlates well with functional sites more evenly, where the highest impact was mediated by select number of residues at the active site (within 5 Å), rather than all of them.

Similarly, we investigated prediction distributions of variation at buried and exposed protein sites using the cytochrome P450 1A1 protein (Uniprot accession: P04798; PDB ID: 6O5Y). In this case, buried residues were those having a surface accessibility of less than 30%, while those which were more than 30% accessible were considered exposed. In general, all three methods predicted buried residues as more pathogenic than their exposed counterparts, a pattern which aligned more notably for AlphaMissense (Figure 4a), and less prominently for VESPA (Figure 4c). This suggests that AlphaMissense more strongly encompasses these 3-dimensional properties, similarly to what was observed for the ATP-binding site in the previous example, despite not being directly trained on this type of data.

Finally, to further assess the usability of these prediction scores on clinical data, we specifically investigated scores from AlphaMissense and ESM1b, which were developed specifically for pathogenicity prediction. In doing so, we assessed the accuracy of these tools, highlighting the proportions of correctly classified variation across both classes, on proteins harbouring at least 10 ClinVar mutations. Notably, both distributions of accuracies overlapped, with AlphaMissense accuracies clustering significantly more at the 95-100% accuracy level, compared to ESM1b (Suppl. Figure 1). This, coupled with the agreement levels summarized in Table 1, highlights comparable capabilities between these two predictors of detecting pathogenic mutations across the human proteome.

## Usage

The two modes of MissenseViewer can be queried using Uniprot^24^ accessions of human proteins, which in the case of “View missense mutations” leads the users to a result page providing access to 3D experimental structures (in RCSB PDB^25,26^) linked to the Uniprot accession, with an additional AlphaFold model covering the complete protein sequence. Each PDB accession is accompanied with the amino acid region resolved in the structure, permitting the user to easily decide which protein to visualise the missense information on (Suppl Figure 2, Section 1). Clicking on a specific structure accession in the table (Suppl Figure 2, Section 2) loads the respective PDB in the structure viewer, permitting colour-coded visualisation of site-specific average AlphaMissense, ESM1b or VESPA scores, so that users can observe general trends in predictions across the protein. The viewer also allows for the creation of publication-quality figures, through the Mol* action menu. Users can select residues of interest by clicking in the sequence viewer at the bottom of the page. This sequence viewer (Suppl Figure 2, Section 3) contains six main parts: i) the reference protein sequence for the Uniprot accession, ii) the imported Interpro-Pfam^27,28^ annotations in the form of sequence coverage, the coverage of the active structure and site-specific susceptibility scores computed from AlphaMissense, v) ESM1b, vi) VESPA and vii) the average of the three predictions. Once a residue is selected, it is highlighted across both sequence and structure viewers, and a table summarising the prediction values for the three predictors is displayed on the right-hand side, where the user can toggle between the three methods. (Suppl Figure 2, Section 4).

Alternatively, in querying “View ligand information” users can assess the predictions of variants within 5 Å of ligands by inputting a Uniprot accession of a protein of interest. A list of all ligands observed in the RCSB PDB structures for the respective protein can be browsed under the ‘Ligand overview’ panel, which summarizes ligand identification codes and more notably, the number of structures describing this ligand protein complex (Suppl Figure 3, Section 1). Once a ligand is selected, two graphing panels are populated, showing the combined distribution of scores from AlphaMissense (AM), ESM1b and VESPA of all variants within 5 Å of the ligand from all structures as continuous distributions (Suppl Figure 3, Section 2) and resolved per structure (Suppl Figure 3, Section 3) with a drop-down menu (Suppl Figure 3, Section 4) allowing users to focus on any of the contributing structures.

## Conclusion

MissenseViewer is an interactive, and easy-to-use resource which permits both 1D and 3D comparisons between AlphaMissense, ESM1b and VESPA pathogenicity predictions. All predictions, including structures and figures are easily downloadable from the web interface (https://biosig.lab.uq.edu.au/missenseviewer/). The integrated API together with the availability of susceptibility scores make the resource invaluable for use with meta predictors. This combined information is expected to help guide more directed therapeutic developments, considering potential pathogenicity hotspots across the human genome.

