## Supplementary material for "On interactive spatial visualisation of pathogenicity predictions": Suppl Figure

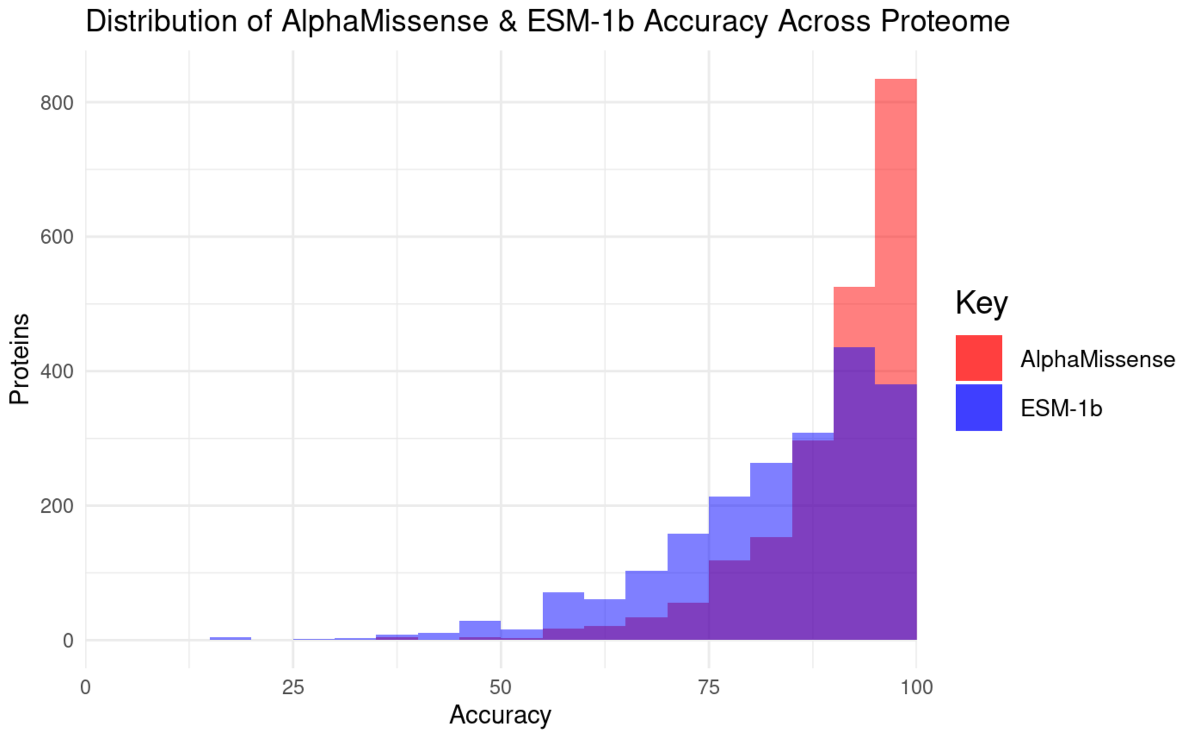


**Suppl. Figure 1: Distribution of AlphaMissense and ESM1b accuracies across proteins harbouring ClinVar mutations.** When considering proteins containing at least 10 ClinVar variants with non-conflicting interpretations, and at least 1-star rating, AlphaMissense and ESM1b had overlapping distributions of accuracies. Notably, AlphaMissense distribution clustered between 95-100% accuracy for over double the number of proteins, compared to ESM1b.

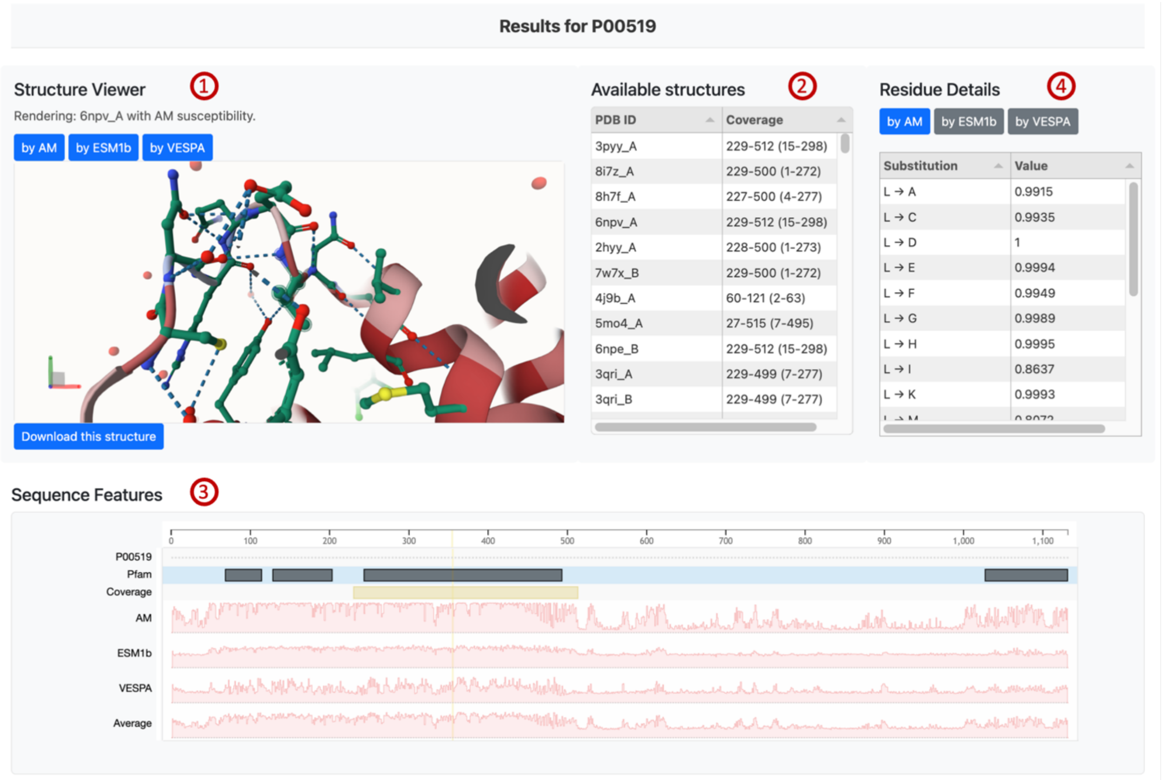


**Suppl. Figure 2: MissenseViewer webserver layout.** Our interactive webserver enables the 3D visualization (1) of proteins of interest, coloured according to AlphaMissense, ESM1b and VESPA scores. Users can choose from a list of structures including the experimentally determined structures in RCSB PDB and the AlphaFold structure (2) and compare their site specific scores from the sequence viewer (3). Clicking on a residue in the sequence viewer, displays AlphaMissense, ESM1b and VESPA predictions (4) at that locus.

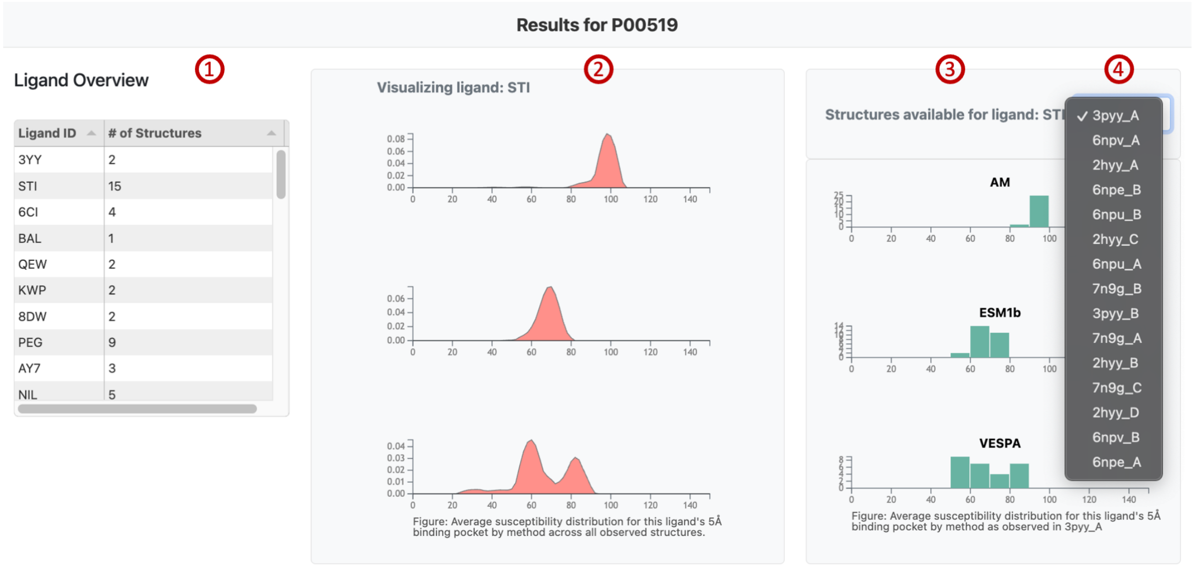


**Suppl. Figure 3: MissenseViewer ligand analysis.** Our interactive webserver also summarises predicted scores of variants within ligand-bound complex structures. A list of ligands (1) binding to the protein of interest, can be browsed, from which, selecting a specific ligand highlights the combined distribution of AlphaMissense, ESM1b and VESPA scores in a continuous (2) or resolved per-structure breakdown (3) format summarising scores of sites within 5 Å. In cases where multiple PDB accessions describe the same protein:ligand complex, the user can choose which accession they are interested in analysing (4).
